# Multicellular PD Control in Microbial Consortia

**DOI:** 10.1101/2023.03.22.533755

**Authors:** Vittoria Martinelli, Davide Salzano, Davide Fiore, Mario di Bernardo

## Abstract

We propose a multicellular implementation of a biomolecular PD feedback controller to regulate gene expression in a microbial consortium. The implementation involves distributing the proportional and derivative control actions between two different cellular populations that can communicate with each other and regulate the output of a third target cellular population. We derive analytical conditions on biological parameters and control gains to adjust the system’s static and dynamical properties. We then validate the strategy’s performance and robustness through extensive *in silico* experiments.

## I. Introduction

The aim of Synthetic Biology is to engineer biological systems with novel functionalities [1]. This is made possible by designing artificial genetic circuits and embedding them into living cells, such as bacteria or yeast, changing their natural behavior to make them, for example, produce some proteins or other chemicals of interest. Applications range from health treatments [2] to bioremediation [3] and production of drugs or biofuels [4]. However, due to the inherently nonlinear and stochastic nature of the biochemical processes involved, reliable and robust regulation of gene expression must be guaranteed by designing synthetic feedback control architectures. Although the control theoretic literature abounds with high performance, robust control solutions, such as MPC algorithms or discontinuous controllers, the need for implementing the control action using biochemical reactions performed inside the cell strongly limits the possible structure of the controllers that can be realized. Hence, simpler feasible solutions have been proposed in the literature (see [5], [6] for an overview); most notably, the antithetic feedback controller realizing an integral action that guarantees robust perfect adaptation with respect to noise and parameters’ uncertainties [7], [8]. Recently, it has been proposed that Proportional-Integral-Derivative (PID) controllers can be implemented via biomolecular reactions and embedded in a single cell to enhance the stability and performance of antithetic controllers, by adding both promptness of regulation response and damping with respect to oscillations [9], [10]. However, in some applications it is more important to avoid oscillations than to ensure a precise regulation, e.g. when engineering the immune response [11]. In these cases a simpler PD control scheme should be preferred because it can guarantee fast and damped regulation of the biological process.

The implementation of biochemical controllers in living cells is also limited by technological and biological constraints, e.g. incompatible chemical reactions, metabolic load or retroactivity [1]. As a consequence, embedding more complex biomolecular controllers consisting of multiple actions (e.g. a PI, PD or PID) inside a single cell can be challenging. A promising solution to these problems is to distribute the control functionalities across different populations in a microbial consortium. The resulting *multicellular* control architecture can both minimize unwanted effects and increase modularity and re-usability of the designed components [1]. In this Letter, we integrate and expand our previous work on multicellular PI controllers [12] by presenting a multicellular PD control strategy where two *controller populations*, each implementing one of the two control actions, regulate a biological process inside a third *target population*, closing the feedback loop by means of diffusing quorum sensing molecules produced by the cells. First, we derive a model of the consortium. Then, we provide some design guidelines for the choice of the control gains based on analytical results, linking the control parameters to the static and dynamical performance of the closed-loop system. The theoretical analysis is complemented with *in silico* experiments carried out via *BSim* [13]. We wish to emphasize that the analysis and design of a fully distributed biomolecular derivative action is a fundamental step together with our previous work in [12] to achieve a fully distributed PID control strategy, which is our ultimate goal.

## II. Multicellular PD control strategy

As stated above, the overarching goal of our design is to engineer a multicellular consortium where two controller populations implement the proportional and the derivative control actions needed to realize a strategy similar to the classical PD control strategy [14],

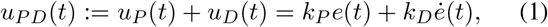

where *k*_*P*_ *>* 0 and *k*_*D*_ *>* 0 are the proportional and derivative gains, *e*(*t*) is the control error defined as the mismatch between the measured output, say *y*(*t*), of the process within a third target population (see Fig. 1) and the desired reference value *Y*_d_(*t*), provided either externally to the controller cells or hard-coded into them as a constitutive promoter. As in [9], here we assume that the desired output, *Y*_d_(*t*) is either a succession of step functions or some constant value.

**Fig. 1:**
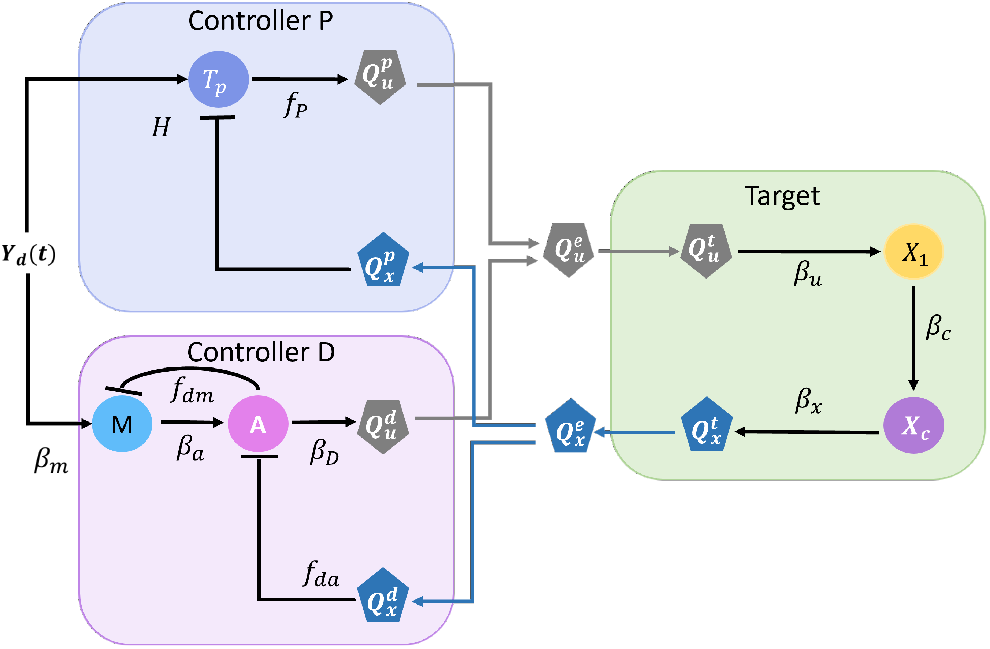
Abstract implementation of the proposed multicellular PD controller. The controller populations compare the reference signal *Y*_d_(*t*) and the quorum sensing molecule *Q*_*x*_, which is produced proportionally to the target gene *X*_*c*_. The process Φ(*t*) is represented by the couple of genes *X*_1_ and *X*_*c*_, in which *X*_1_ is the actuated by the quorum sensing molecule *Q*_*u*_. Circles represent internal molecular species, while polygons are the signaling molecules.

The abstract biological implementation of our proposed distributed PD control strategy is shown in Fig. 1. Therein, each controller population computes the control error *e*(*t*) by comparing the reference signal *Y*_d_(*t*) with the measure of the output *y*(*t*) corresponding to the expression level of *X*_*c*_ within the target population. Such measure is broadcast to the whole consortium via the quorum sensing molecule *Q*_*x*_ which acts as a proxy for *y*(*t*). The control input *u*_*PD*_(*t*) computed by the two controller populations is delivered to the target cells by means of the quorum sensing molecule *Q*_*u*_, that is produced by the controllers and effectively serves as the “actuating” signal in this multicellular control scheme. To avoid cross-talking effects [15], *Q*_*x*_ and *Q*_*u*_ are assumed to be orthogonal.

We will show next that the reactions depicted in Fig. 1 implement a distributed biomolecular PD controller.

### A. Mathematical Modelling

We derive a model capturing the *aggregate* dynamics of the microbial consortium, that is the evolution of the concentrations of the biochemical species averaged over all cells in a population. (Details on the derivation of the aggregate dynamics from the single cell dynamics are reported in Appendix C.)

As also done in [12], we make the following simplifying assumptions: (i) all populations in the consortium are equally balanced, that is, the three populations are composed by the same number of cells, *N* ; (ii) the total number of cells in the consortium is constant. Note that the former assumption can be achieved by means of external ratiometric controllers, e.g. [16], while the latter assumption is generally verified in experimental environments with limited space and resources, e.g., microfluidics, chemostats, vials. From these assumptions it follows that the diffusion dynamics of the quorum sensing molecules through the cells’ membranes can be supposed to have the same diffusion rate *η*. (This assumption will be relaxed later in Section IV). In what follows, the superscripts *e, t, p, d*, are used to denote quantities in the “environment”, in the “target” cells, in the “proportional”, or in the “derivative” controller cells, respectively.

#### 1) Target cells

As done in [7], [9], we assume that the biological process to be controlled inside the target cells consists of only two genes, *X*_1_ and *X*_*c*_ (Fig. 1). The target gene *X*_*c*_ is directly activated by *X*_1_, which in turns is actuated by the quorum sensing molecule *Q*_*u*_ produced by the controller cells. Using mass-action kinetics, the dynamics of the genetic network inside the target cells can therefore be described by the following set of ODEs:

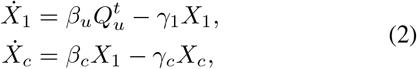

where *β*_*u*_ and *β*_*c*_ are activation rates describing the strength of the transcription factors *X*_1_ and *Q*_*u*_, while *γ*_1_ and *γ*_*c*_ are degradation rates.

The output of the process, corresponding to the expression level of *X*_*c*_, is broadcast to the other cells by means of the diffusing quorum sensing molecule *Q*_*x*_, which is produced by the target cells. Assuming as in [12] that the production of *Q*_*x*_ is proportional to *X*_*c*_, the dynamics of the sensing molecule inside the target cells is described by:

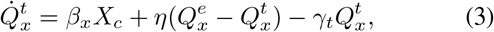

where *β*_*x*_ is the activation rate of *Q*_*x*_, *η* is the diffusion rate of the molecule across the cell membrane, and *γ*_*t*_ is the dilution rate into the target cells.

#### 2) Proportional controller cells

The Proportional controller is implemented here as in [9] as a nonlinear function *f*_*P*_ = *f*_*P*_ (*Q*_*x*_, *Y*_d_), activating the production of *Q*_*u*_. Specifically, the dynamics of the concentration of *Q*_*u*_ in the Proportional controllers is given by:

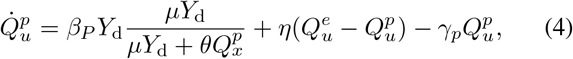

where *γ*_*p*_ is the dilution rate of the quorum sensing molecules into the Proportional cells, *μ* and *θ* are positive control parameters, and *β*_*P*_ is a tunable parameter which plays the role of a proportional gain. It has been shown in [9] that the first term in (4) realizes a control action that is a function of the control error *e*(*t*) := *μY*_d_ *−θQ*_*x*_.

#### 3) Derivative controller cells

An approximation of the time derivative of the control error *e*(*t*) can be obtained by embedding in the controller population the biomolecular circuit represented in Fig. 1 (see Appendix A for further details), whose dynamics can be described as follows:

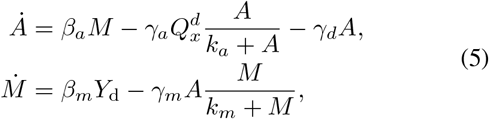

where *γ*_*d*_ is the dilution rate of *A* due to cell growth and division, and *β*_*a*_ and *β*_*m*_ are activation rates, which in turns are actively degraded by *Q*_*x*_ and *A*, respectively, through enzymatic reactions modeled here via Michaelis-Menten functions with constants *γ*_*a*_, *k*_*a*_, *γ*_*m*_, *k*_*m*_. For the sake of simplicity, we assumed here that the quorum sensing molecule *Q*_*x*_ also directly acts as a degradation enzyme on *A*. The derivative control action is then delivered to the targets by means of the quorum sensing molecule *Q*_*u*_, produced by the controllers at a rate proportional to *A* (and thus proportional to an approximation of *ė*(*t*)), whose concentration inside the controllers is given by:

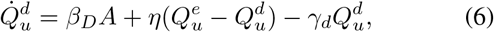

where *β*_*D*_ plays the role of a derivative gain, and the other parameters have an analogous meaning to those in (3).

In addition, the dynamics of *Q*_*x*_ and *Q*_*u*_ into cells where they are not produced is described by:

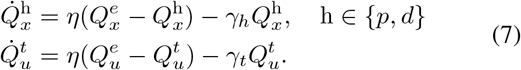

Finally, the model is completed by describing the evolution of the concentrations of the quorum sensing molecules into the environment:

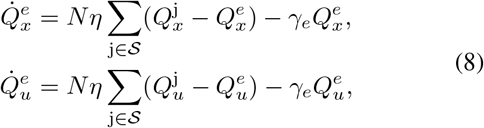

where *N* is the number of cells in each population (assumed to be balanced), and 𝒮 = {*p, d, t*}.

## III. Design of the microbial consortium

The goal of the multicellular PD architecture shown in Fig. 1 and modelled by (2)-(8) is to regulate the concentration of the quorum sensing molecule 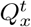, that acts as a proxy of the target output *X*_*c*_, to the set-point specified via the reference signal *Y*_d_ with given static and dynamic performance. Specifically, we assume the desired maximum amplitude of the steady state error to be *ϵ*_*e*_ with a settling time less than *ϵ*_*t*_ and an overshoot *s*_%_ ≤*ϵ*_*s*_. Note that these bounds may vary depending on the specific application of interest. Hence, in this work we will not fix these quantities *a priori*. Instead, we will investigate how the control gains can be chosen so as to modulate the settling time, the overshoot and the steady state error that can be achieved.

We will show that, as for the classical PD control, the addition of the derivative action to the consortium can provide a prompter and better regulation of the process output inside the target cells. However, the control gains *β*_*P*_ and *β*_*D*_ in (4) and (6) need to be appropriately tuned to achieve the desired stability and performance. Next, we provide analytical conditions on the control gains that can be used to tune the static precision and the characteristics of the transient response of the closed-loop system, and we compare them to those of the simpler Proportional controller presented in [12].

## A. Reduced order model

To derive meaningful analytical conditions we make the following simplifying assumptions, commonly made in the literature, e.g., [9], [12], [17].

**Assumptions** *The multicellular control system* (2)-(8) *satisfies the following proprieties:*

A1 *All cell populations grow and divide at the same rate*.

A2 *The quorum sensing molecules Q*_*u*_ *and Q*_*x*_ *diffuse faster than they degrade*.

A3 *The enzymatic reactions in* (5) *occur with saturation of the substrates*.

A4 *The dynamics of the derivative action* (5) *is sufficiently faster than the controlled process in* (2).

### Remark 1

*Assumption A1 implies that all cells have the same dilution rate, i*.*e. γ*_1_ = *γ*_*c*_ = *γ*_*p*_ = *γ*_*d*_ = *γ*_*t*_ = *γ, which is a reasonable assumption if all populations are realized using the same biological chassis, and therefore, despite inevitable fluctuations, grow and divide at the same rate. Assumption A2 implies that η* ≫Γ_*PD*_, *with* Γ_*PD*_ := *γ*_*p*_ + *γ*_*d*_ + *γ*_*t*_ + *γ*_*e*_ = 3*γ* + *γ*_*e*_, *and it is common in the literature and holds in different models parameterized from in vivo experiments, see e*.*g. [17]. Assumption A3 is commonly made in the literature (see for example [18]) and it implies that K*_*a*_ ≪*A and K*_*m*_≪ *M [19]. Assumption A4 allows to perform a time scale separation between the dynamics of the process and the one of the derivative controller, and it can be satisfied by requiring that the frequency content of X*_*c*_(*t*) *(and thus of Q*_*x*_(*t*)*) is bounded by some value ω*_max_, *and the parameters of the derivative motif are chosen accordingly (see [9] and Appendix A for details)*.

Under the previous assumptions, the dynamics of the multicellular system can be approximated by the following reduced order model (see Appendix A for details):

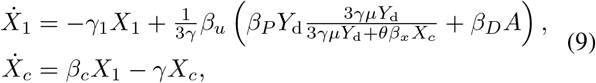

with *A* defined as in (18). Note that the transfer function of system (9) can be locally mapped to the equilibrium point of the closed-loop system to the one of a classic PD controller, see [9] for further details.

### B. Static and transient performance

Setting 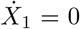 and 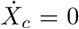 in (9), the equilibria can be obtained as

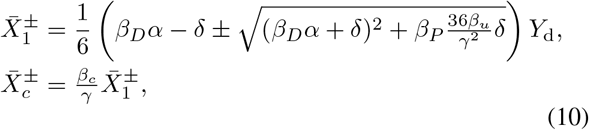

where 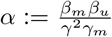 and 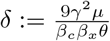. Since the state variables *X*_1_, *X*_*c*_ describe the concentration of chemical species, we restricted the admissible solutions to be positive. And thus, since 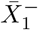 is negative for any positive values of the system parameters, the only admissible equilibrium point is 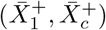, which, by applying the Routh-Hurwitz criterion, can be proved to be locally asymptotically stable for any value of the parameters (see Appendix D for details on the stability analysis).

Next, from the expression of the control error at steadystate (the derivation is omitted here for the sake of brevity), we find that if the control gains are positive (*β*_*P*_, *β*_*D*_≥ 0) and such that:

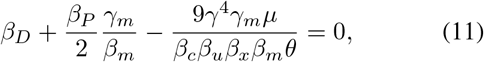

the steady-state error can theoretically be made null provided that all parameters are perfectly known. In practice, this is unrealistic in biological implementations of the controller and therefore the error can only be made small enough by appropriately tuning the gains around the values suggested in (11).

We assess the (local) transient performance by studying the eigenvalues of the linearization of system (9) when (11) is fulfilled about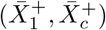, where, substituting (11) into (10), 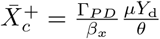. Simple algebraic manipulations yield:

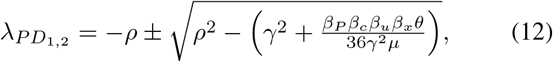

where 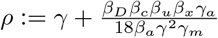.

The eigenvalues are therefore both real and negative if:

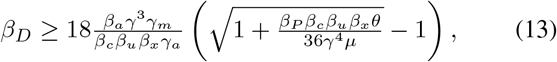

and correspondingly the transient response is characterized by the absence of oscillations, otherwise they are complex conjugates with negative real part.

### C. Comparison with the Proportional controller

To highlight the advantages of adding a derivative control action to the multicellular control scheme, we compare the static and transient performances of the PD controller with those of the P controller alone we previously studied in [12]. Regarding the steady-state, also in the case when only a proportional biomolecular controller is present, the error can be made theoretically null (and therefore closer to zero in practice) provided that all the system parameters are perfectly known and the proportional gain is selected about the value [12]:

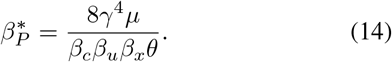

Following similar steps as those taken to obtain (12), we find that in the absence of the derivative action when *β*_*P*_ is set as in (14), the transient response is governed by the pair of complex conjugate eigenvalues:

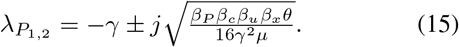

Thus, when the derivative action is added to the consortium, if the gains are tuned so that 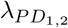 are real, i.e. condition (13) holds, by comparing (12) with (15) we find that the PD strategy can indeed guarantee faster convergence (that is, 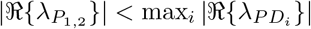 by further requiring that:

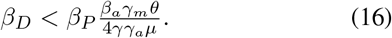

Also, if 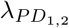 are complex conjugate, it is easy to verify that *ℜ* {*λ*_*PD*_} *< ℜ* {*λ*_*P*_} and *ℜ* {*λ*_*PD*_} *< ℜ* {*λ*_*P*_} for any choice of the control gains, therefore, the PD controller always guarantees faster response and more damped oscillations than the Proportional controller alone.

## IV. In silico Experiments

We validated the proposed PD multicellular architecture via *in silico* experiments carried out in Matlab and BSim [13] using the model described by equations (2)-(8).

First, we validated the theoretical predictions using Matlab. Specifically, as reported in Fig. 2, we computed for different values of the gains *β*_*P*_ and *β*_*D*_ the overshoot *s*_%_, the 5% settling time 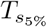, and the relative percentage steadystate error defined as 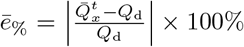, where 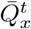 is the steady-state value of 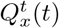 and 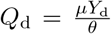. We found that, as expected from the analysis, the PD strategy can guarantee bounded steady-state error for gains chosen close to the theoretical estimates reported in (11) (*β*_*D*_ *<* 0.17, *β*_*P*_ = 0.09 *−*0.54 *β*_*D*_). For the sake of completeness we also report in Fig. 3 the steady-state error, overshoot and settling time when only the Proportional control action is present (*β*_*D*_ = 0). Also we notice that, as predicted by (12) and (15), the proportional action used on its own always causes overshoots in the closed-loop response whereas there is a wide range of control parameters (see Fig. 2.b) where the PD action ensures no overshoot. Also, when condition (16) holds (*β*_*D*_ *<* 4.89 *β*_*P*_), the PD architecture provides faster dynamic regulation than the Proportional one (see Fig. 2.c). We further validated our theoretical results by carrying out agent-based simulations in BSim accounting for cell growth, cell-to-cell variability, diffusion and the geometry of cells and of the hosting chamber. In all the simulations, we set the initial concentration for all chemical species to zero. This emulates an experimental protocol where each cellular population is grown separately from the others prior to the beginning of the experiment. We assumed the cells are growing in a scaled-down version of the microfluidic chamber used in [16], [20]. Specifically, the chamber has dimensions 17*μ* m *×*15*μ* m *×*1 *μ*m and can host around 100 cells, which is a good compromise between computational time and statistical relevance. Unless otherwise stated, the growth and mechanical parameters of cells and the nominal values of the parameters in the network were chosen as described in Appendix B.

**Fig. 2:**
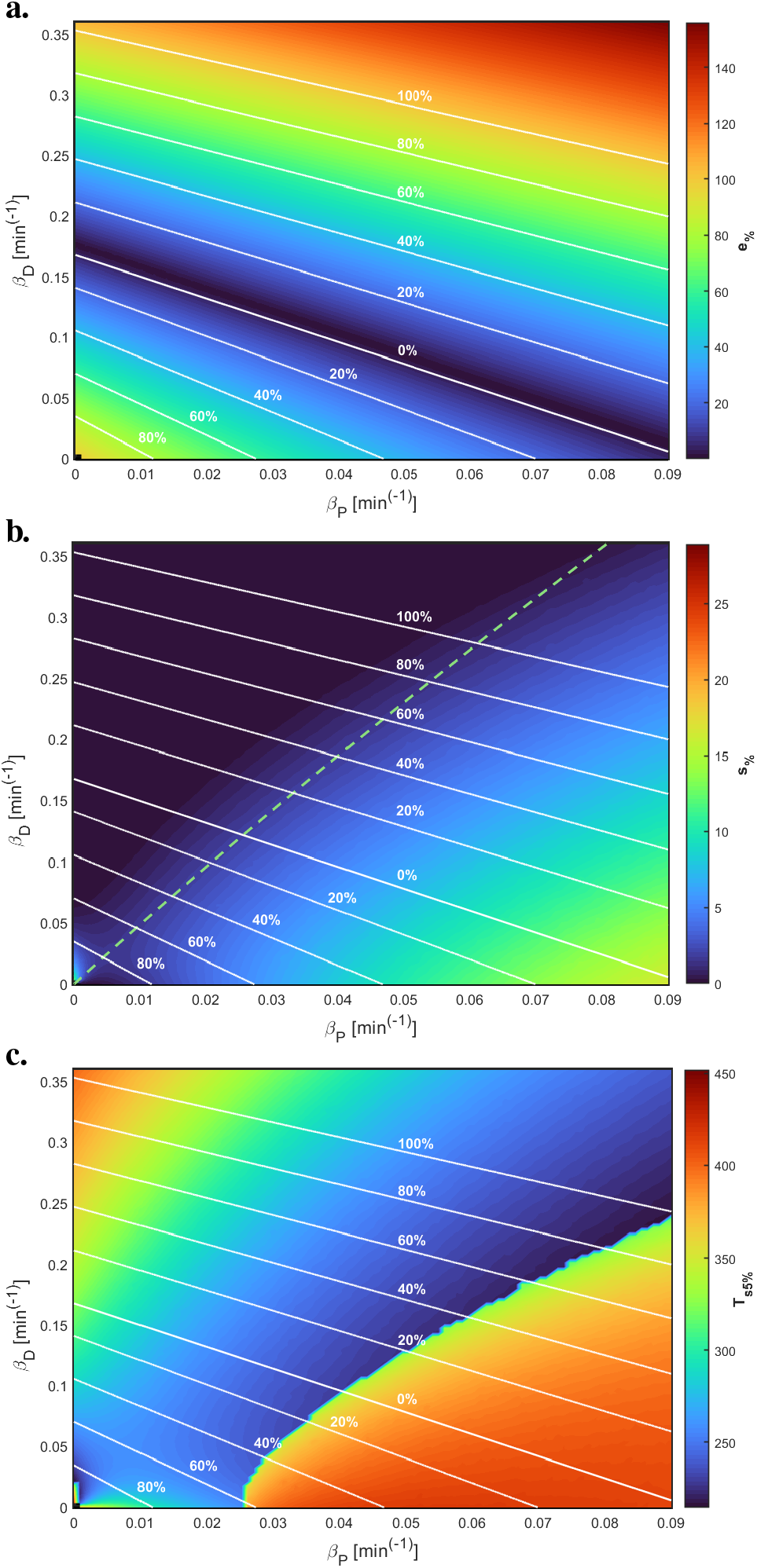
*In silico* experiments in Matlab: percentage error *e* (a), percentage overshoot *s*_%_(b) and settling time 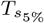 (c) when the targets are controlled by the PD controller. The control gains *β*_*P*_ and *β*_*D*_ were varied in the intervals [0, 0.09] and [0, 0.36] with step size equal to 0.0009 min^*−*1^ and 0.0036 min^*−*1^, respectively. The reference signal was set to *Y*_d_ = 60 nM in all simulations. The white isoclines bound the regions in the parameters’ space (*β*_*P*_, *β*_*D*_) with the same upper bound on the percentage error. The 0% isocline in panel (a) is the analytical estimate given by (11) while the green dashed line in panel(b) represents the condition to have real eigenvalues given by equation (13).

**Fig. 3:**
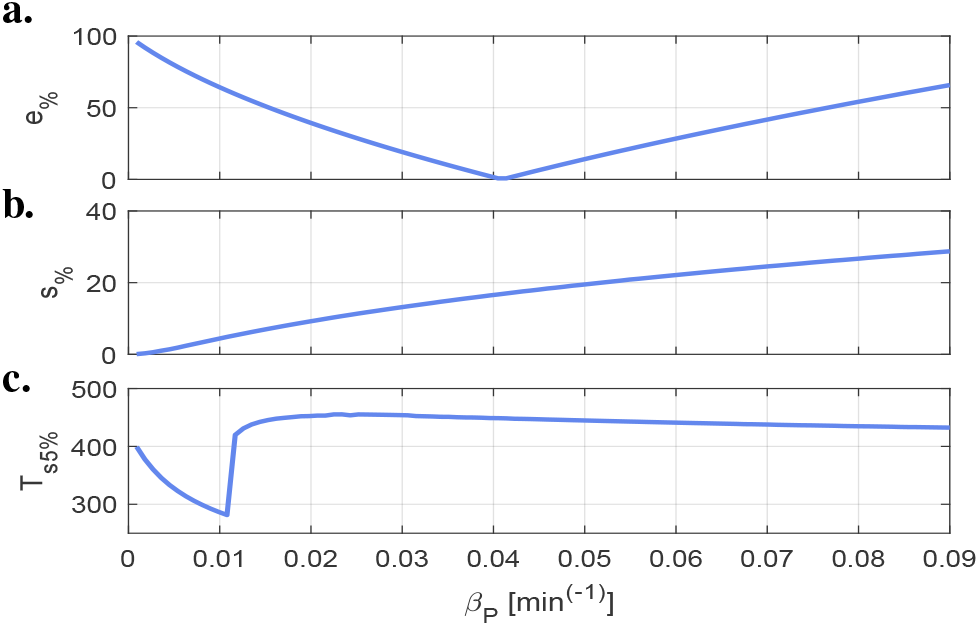
Proportional performance validation in Matlab: percentage error *e*_%_ (a), percentage overshoot *s*_%_ (b) and settling time 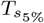 (c) when the targets are controlled by a proportional controller. The control gain was varied in the interval [0, 0.09] with step size equal to 0.0009 min^*−*1^. The reference signal was set to *Y*_d_ = 60 nM in all simulations.

We chose values of the control gains *β*_*P*_ and *β*_*D*_ that were shown in Matlab (see Fig. 2) to guarantee *e*_*∞*_ *<* 20% when *Y*_d_ = 60 nM and to also satisfy condition (16). We ran 10 agent-based simulations, each time randomly selecting the control gains with uniform distribution and evaluating the performance. Figure 4 shows that both the P and PD architectures have good regulation capabilities. Moreover, it confirms that on average the addition of a derivative action consistently reduces the overshoot (9.76% for the PD, and 18.29% for the P controller). Instead, the reduction of the settling time was just about 10% (about 355 min for the PD, and 392 min for the Proportional). As for the steady-state error (4.55% for the PD, and 2.71% for the Proportional), the difference between the two strategies was negligible.

**Fig. 4:**
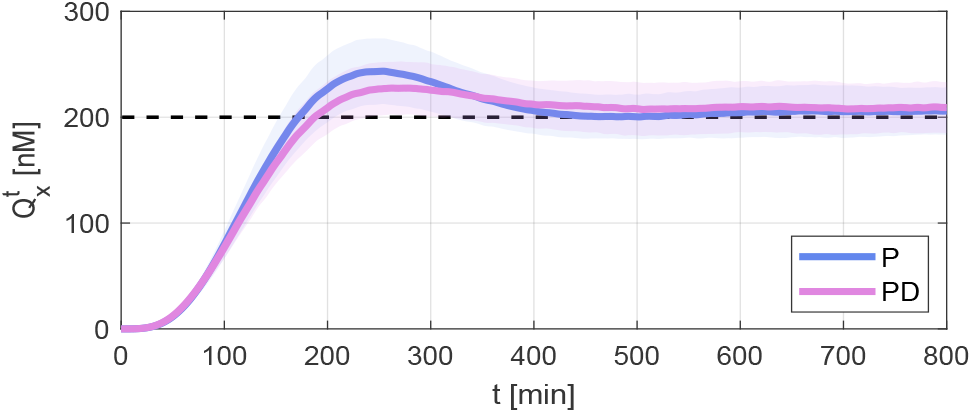
Control performance validation in BSim: mean (continuous line) and standard deviation (shaded area) of the average concentration of 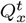 in the targets. The statistics were computed over 10 simulations. The blue lines are obtained using a Proportional controller and the purple ones using a PD controller. Control gains were drawn with uniform distribution from the region of the parameter space (*β*_*P*_, *β*_*D*_) in Figure 2 in which *ē*_%_ *<* 20%. For the PD architecture we also imposed condition (16). The reference signal *Q*_d_ = *μY*_d_*/θ* is depicted as a dashed line. At *t* = 0 min, 18 cells, equally divided between the populations, were positioned at the center of the chamber along a horizontal stripe.

Finally, we tested the robustness of the P and PD strategies as the cell-to-cell variability increases in the cell populations. We modeled this effect by drawing, at cell division, each parameter of the daughter cells, say *ρ*, from a normal distribution centered at its nominal value 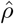 with standard deviation 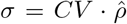, where *CV* is the coefficient of variation. We compared the robustness of the two architectures, as the *CV* increases, by evaluating the settling time, the overshoot, and the average relative percentage steady-state error, defined as 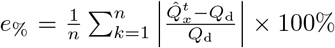, where 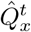 is the value of 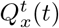, averaged over the last 200 min, of the *k*-th experiment, 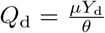, and *n* is the total number of experiments we conducted for each value of *CV*. Fig. 5 shows that the P and the PD strategies both possess good robustness (the steady-state error never exceeds 20%). We observe that while the Proportional controller guarantees a lower residual error under perturbations (Fig. 5.a), the PD control strategy reduces the overshoot even in the presence of high cell-to-cell variability (Fig. 5.c); no significant difference between the two being detected in terms of settling time (Fig. 5.b).

**Fig. 5:**
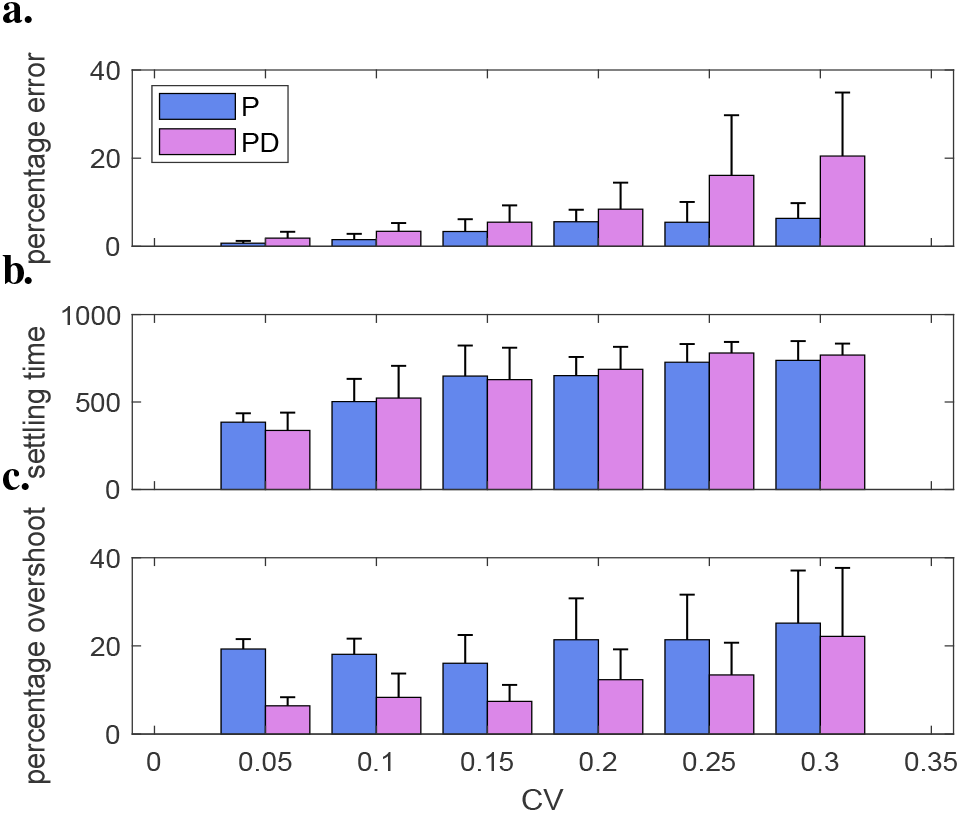
Sensitivity to parameter variations: mean and standard deviation of the percentage of error at steady-state (a), settling time (b) and overshoot (c) as the cell-to-cell variability increases. Blue bars are obtained using only a Proportional controller and purple bars when a PD controller was employed. For each value of *CV ∈* {0.05, 0.1, 0.15, 0.2, 0.25, 0.3} we performed *n* = 50 simulations drawing independently cells’ parameters from a normal distribution centered at their nominal value 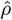 with standard deviation 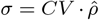. The reference signal was set to *Y*_d_ = 60 nM, while the gains were chosen as *β*_*P*_ = 0.0414 min^*−*1^ and *β*_*D*_ = 0.0933 min^*−*1^. All simulations were performed for a total time of 800 min.

## V. Conclusions

We discussed the implementation of a distributed biomolecular PD controller. Analytical conditions were derived for control gains to achieve the desired performance. It was shown that the derivative action helps reduce overshoots and settling time. Numerical and *in silico* experiments were conducted to confirm the effectiveness of the proposed strategy, despite cell-to-cell variability and other realistic effects. Future work involves exploiting the results presented here with our previous results in [12] to achieve a fully distributed biomolecular PID controller.

## Appendix

### A. Derivation of the reduced order model (9)

Under Assumption A2, we can assume that the quorum sensing molecules are always at steady state. Therefore, by imposing that 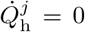 and 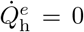, with h *∈* {*u, x*} and *j∈ 𝒮*, where 𝒮 = {*t, p, d*}, and under the hypothesis of a large number of cells, after some algebra we have that:

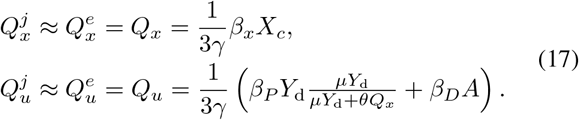

Under Assumption A4 there exists some *ω*_max_ upper bounding the frequency contents of *Q*_*x*_(*t*), therefore, if the parameters of the derivative motif are chosen so as to satisfy the condition 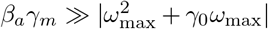, then *A* in (5) contains an approximation of the derivative of the control error *e*(*t*) given by (see [9] for further details):

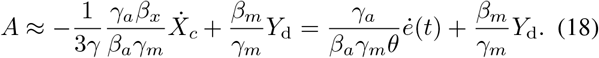

This is because, under Assumption A2 we can substitute the expression of *Q*_*x*_ at steady-state in (17) in the control error *e*(*t*). Then, assuming a constant reference signal *Y*_d_ and taking the time derivative of *e*(*t*), yields:

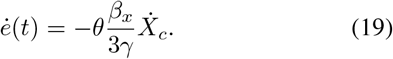

Combining the quorum sensing steady-state expressions in (17) and (18) with (2) we then obtain the reduced model (9).

### B. Nominal biochemical parameters

The growth and mechanical parameters used in the BSim simulations were selected as in [17], while the nominal biochemical parameters were chosen as: *β*_*u*_ = 0.06 min^*−*1^, *β*_*x*_ = 0.03 min^*−*1^, *γ* = 0.023 min^*−*1^, *η* = 2 min^*−*1^ (taken from [17]); *β*_*c*_ = 0.1 min^*−*1^, *μ* = 1 min^*−*1^, *θ* = 0.3 min^*−*1^, *β*_*a*_ = 1.5 min^*−*1^, *β*_*m*_ = 0.4167 min^*−*1^, *γ*_*a*_ = *γ*_*m*_ = 1.5 min^*−*1^ (taken from [9]); *γ*_*e*_ = 0.0023 min^*−*1^.

### C. Derivation of the aggregate model

Here we present the derivation of the aggregate model reported in Section II-A that describes the evolution of the average concentrations of chemical species in the whole consortium. For the sake of brevity, we report there only the case of a consortium comprising Proportional controller cells and target cells. The case of three populations, in which also the derivative controller population is present, follows straightforwardly. The aggregate dynamics of the non-diffusive species (i.e., *X*_1_ and *X*_*c*_ in the target cells, and *A* and *M* in the derivative controller cells) follows directly from the single cell model by assuming that the average concentrations follow the same dynamics. For what concerns the diffusive species (i.e., *Q*_*x*_ and *Q*_*u*_), under the assumptions made in Section III-A, their dynamics inside each cell in the consortium and into the environment is given as (we report here only the dynamics of *Q*_*x*_ for the sake of brevity, a similar analysis follows for *Q*_*u*_):

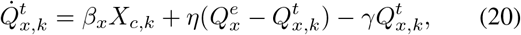

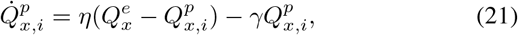

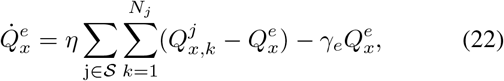

where *X*_*c,k*_ and 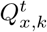 are the concentrations of the species *X*_*c*_ and *Q*_*x*_ inside the *k*-th cell of the target population, 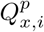 is the concentration of the signaling molecule *Q*_*x*_ in the *i*-th cell embedding the Proportional controller, and 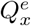 is the concentration of *Q*_*x*_ into the environment. Moreover, 𝒮 = {*t, p*}, and *N*_*t*_ and *N*_*p*_ are the number of cells in the target and Proportional controller populations, respectively. Now, defining 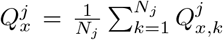 for *j ∈ 𝒮* and 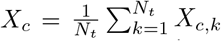, we can derive the evolution of the average concentration of *Q*_*x*_ inside each cell population as:

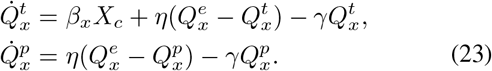

In addition, we can recast equation (22) as:

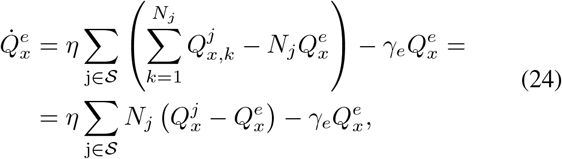

and thus, assuming *N*_*t*_ = *N*_*p*_ = *N*, that is, the two populations are balanced, equation (24) becomes:

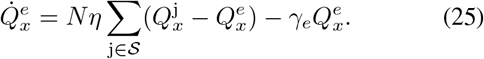

### D. Stability analysis of the PD control model

The local stability of system (9) can be assessed by computing the characteristic polynomial linearized about the admissible equilibrium point 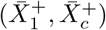 (10), and applying the Routh-Hurwitz criterion [21]. For a second order system the Routh table is given by:

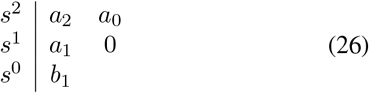

The elements in table (26) are computed starting from the coefficients of the characteristic polynomial:

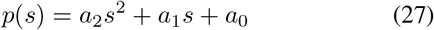

For the system in exam, they can be formalized as

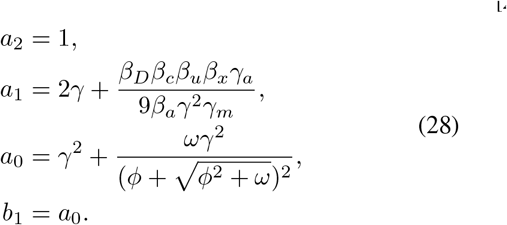

where *ω* := 36*β*_*P*_ *β*_*c*_*β*_*u*_*β*_*x*_*γ*^4^*γ*^2^ *μθ* and *ϕ* := 9*γ*^4^*γ*_*m*_*μ* + *β*_*D*_*β*_*c*_*β*_*u*_*β*_*x*_*β*_*m*_*θ*. Since all the parameters of the system are assumed to be positive constants, all the coefficients in equation (28) are positive. As a consequence, no sign permutations happen in the first column of (26) and, applying the Routh-Hurwitz criterion, we can infer that the equilibrium point is locally asymptotically stable.

